# Genetic influences vary by age and sex: Trajectories of the association of cholinergic system variants and theta band event related oscillations

**DOI:** 10.1101/2023.02.27.530318

**Authors:** David B. Chorlian, Jacquelyn L. Meyers, Niklas Manz, Jian Zhang, Chella Kamarajan, Ashwini Pandey, Jen-Chyong Wang, Martin Plawecki, Howard Edenberg, Alison Goate, Jay Tischfield, Bernice Porjesz

**Affiliations:** SUNY Downstate HSU; College of Wooster; Mount Sinai School of Medicine; Indiana University; Rutgers University

## Abstract

To characterize systemic changes in genetic effects on brain development, the age variation of the associations of cholinergic genetic variants and theta band event-related oscillations (EROs) was studied in a sample of 2140 adolescents and young adults, ages 12 to 25 from the COGA prospective study. The theta band EROs were elicited in visual and auditory oddball (target detection) tasks and measured by EEG recording. Associations were found to vary with age, sex, task modality (auditory or visual), and scalp locality. Seven of the twenty-one muscarinic and nicotinic cholinergic SNPs studied in the analysis, from *CHRM2, CHRNA3, CHRNA5*, and *CHRNB4*, had significant effects on theta band EROs with considerable age spans for some sex-modality combination. No SNP-age-modality combination had significant effects in the same direction for males and females. Results suggest that nicotinic receptor associations are stronger before age 18, while muscarinic receptor associations are stronger after age 18.

## 1 Introduction

The age variation in adolescents and young adults of genotypic effects on theta band event-related oscillations (EROs) was determined by the assessment of developmental trajectories of theta band (4-7 Hz) EROs by non-parametric regression in a sample of 2140 adolescents and young adults ages 12 to 25 from the COGA prospective study, a multisite collaboration designed to study the genetics of alcoholism (Begleiter et al., 1995). The theta band EROs occurring in the P3 response, important indicators of neurocognitive function, were elicited during the evaluation of task-relevant target stimuli in visual and auditory oddball tasks from 9 electrode locations. These tasks call upon attentional and working memory resources. The developmental trajectories of the EROs have large sex differences; scalp location and task modality (visual or auditory) differences within males and females were small compared to sex differences. Associations between the theta EROs and genotypic variants of 21 single nucleotide polymorphisms (SNPs) from cholinergic genes *CHRM2, CHRNA3, CHRNA5, CHRNA6, CHRNB1, CHRNB3*, and *CHRNB4*, measured by effect sizes of genotypic covariates in the regression analysis, were found to vary with age, sex, task modality, and scalp locality. Seven of the SNPs had significant effects with considerable age spans for some sex-modality combination. Eight of the SNPs had significantly different effects in males and females with considerable age spans for either one or both modalities. No SNP at any age-modality combination had significant effects in the same direction for males and females. Principal component analysis (PCA) of the genotypic profiles (vector of values of genotypic associations) of phenotypes identified the age variation in genotypic profiles which discriminate sex, modality, and locality differences in phenotypes. The physiological processes responsible for large sex differences in genotypic effect on brain function have yet to be determined.

### Prefatory Note

The contents of this paper were taken directly from a poster presentation at the Human Genenetics in New York Conference of February 2, 2017. Supplementary material derived from documents which were antecedents of the presentation text is found in section 8.

## 2 Theta EROs: Phenotypic and Association Trajectories

### 2.1 Background and Methods

In a previous study (Chorlian et al., 2015), developmental trajectories of theta EROs extending from ages 12 to 25 were determined by non-parametric regression (loess) using as covariates sex, family type (control or alcoholic), and the first two principal components from the stratification analysis of the genetic dataset. Following the determination of the trajectories of the ERO values the trajectories of the genotypic associations of the ERO values were determined. The developmental trajectories (time-series) of the association between cholinergic gene variants and theta band EROs are defined by the effect size of the genetic variants in the non-parametric regression model of the theta band EROs, to which genotype (using an additive model) and a genotype × sex interaction, were added as covariates. For this study, the association of theta band EROs at nine scalp locations in two experimental modalities and 21 SNPs at 131 age centers (ages 12 through 25 at 0.1 year intervals) were calculated. This provides effect size matrices of size 36 × 21 for each age center. The methods used to obtain the results are described in detail in Chorlian et al. (2017) and in Section 8.2. Much of the analysis was developed as an adaptation of the methods of a gene expression study of brain development in rats by Stead and colleagues (Stead et al., 2006). The key analogy is between the level of gene expression and the effect size of genotypic variants; more details can be found in 8.1.

### 2.2 Phenotypic trajectories

Illustrations of theta ERO trajectories from the central electrodes are shown from Chorlian et al. (2015).

As can be seen in Figure 1, large differences in developmental trajectories between males and females were found; scalp location and task modality differences within males and females were smaller than the sex differences. The figure illustrates both the mean ERO values across the age range (top row) and the rates of change of these values (bottom row) (Chorlian et al., 2015, Section 3.1). Variation in phenotypic correlations (Chorlian et al., 2015, Figure 2) suggests that the phenotypes fall into four groups distinguished by sex and modality.

**Figure 1:**
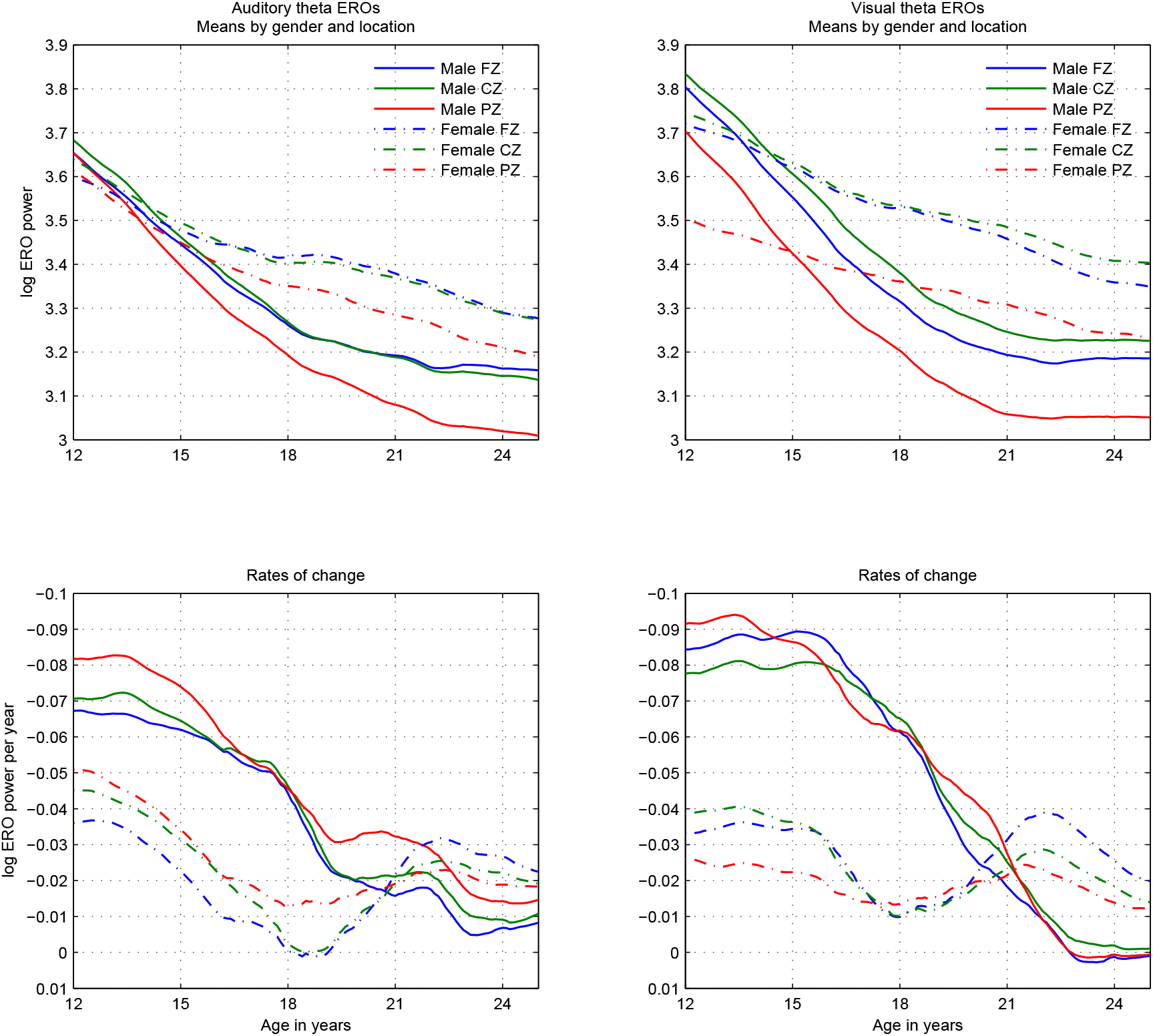
Theta ERO trajectories. Theta ERO total power trajectory means in auditory (left column) and visual (right column) modalities presented in two views: **Top row:** Development curves; **Bottom row:** Rates of change of development curves: Each line in this graph represents the slope of the corresponding line in the graph in the top row at the corresponding age. The y-axis is inverted in order to more clearly illustrate the decrease in absolute value of the slopes with time. Line styles and colors of the bottom graph follow the legend in the top row.

**Figure 2:**
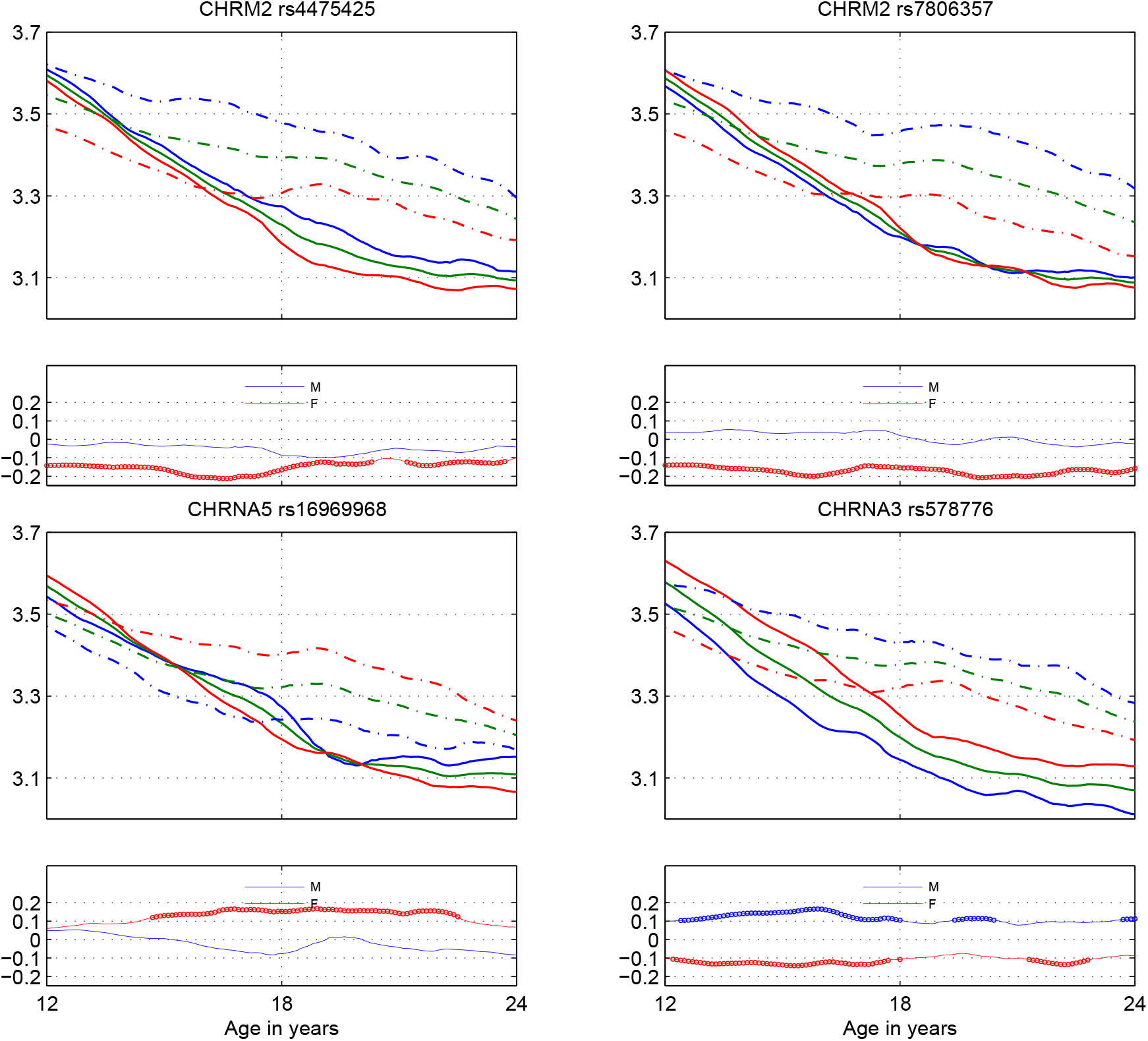
Examples of Association Trajectories: Rows 1 and 3: Genotypic effects of Cholinergic SNPs on Frontal Auditory ERO trajectories from regression model. Solid: Males, Dashed: Females; Blue: 0 Major Alleles, Green: 1 Major Allele, Red: 2 Major Alleles. Rows 2 and 4: Genotypic association effect size (of major allele) trajectories: Blue: Male; Red: Female. Thicker lines indicate significant effect sizes, which correspond to greater inter-allelic distances in Rows 1 and 3

### 2.3 Association trajectories

Illustrations of association trajectories for frontal auditory theta ERO with two muscarinic receptor SNPs and two nico-tinic receptor SNPs are shown as calculated in this study.

As examples of association trajectories, Figure 2 shows genotypic effects of 4 cholinergic SNPs on frontal auditory ERO trajectories from the regression model in rows 1 and 3, shows how mean phenotypic trajectories vary by genotype. These lines should be compared to the blue lines (FZ electrode) in the top left panel of Figure 1 to see the allelic split compared to the overall mean. The corresponding association trajectories of the SNPs, expressed in standard deviations of the data, are shown in rows 2 and 4. The large separation between the lines in rows 1 and 3 corresponds to the enhanced colors in rows 2 and 4. Note the opposite effect sizes for males and females for the nicotinic receptor SNPs, particularly the CHRNA3 SNP.

## 3 Cholinergic - Theta ERO Associations: SNP specific results

### 3.1 Methods

Significant SNPs and significant sex differences for the sex-modality-age combinations were determined by setting threshold effect sizes and *p*-values using a coordination of control of the false discovery rate by the method in Storey and Tibshirani (2003) using *p*-values (See Section 8.3.1 for details), and permutation tests on the absolute values of the effect sizes to establish SNP specificity of results (See Section 8.3.2 for details). Analysis of the distribution of *p*-values establishes a false discovery rate of less than 5% for a threshold of *p* < 10^− 4^. Individual effect sizes were aggregated over age ranges and the nine electrode array in order to minimize effects of spurious large effect sizes on the identification of significant SNPs.

### 3.2 Results

Results are illustrated for 7 overlapping 3 year age ranges extending from age 12 to 24.

Figure 3 shows significant SNPs in the auditory modality. Note that no SNP-age combination is simultaneously significant for males and females.

**Figure 3:**
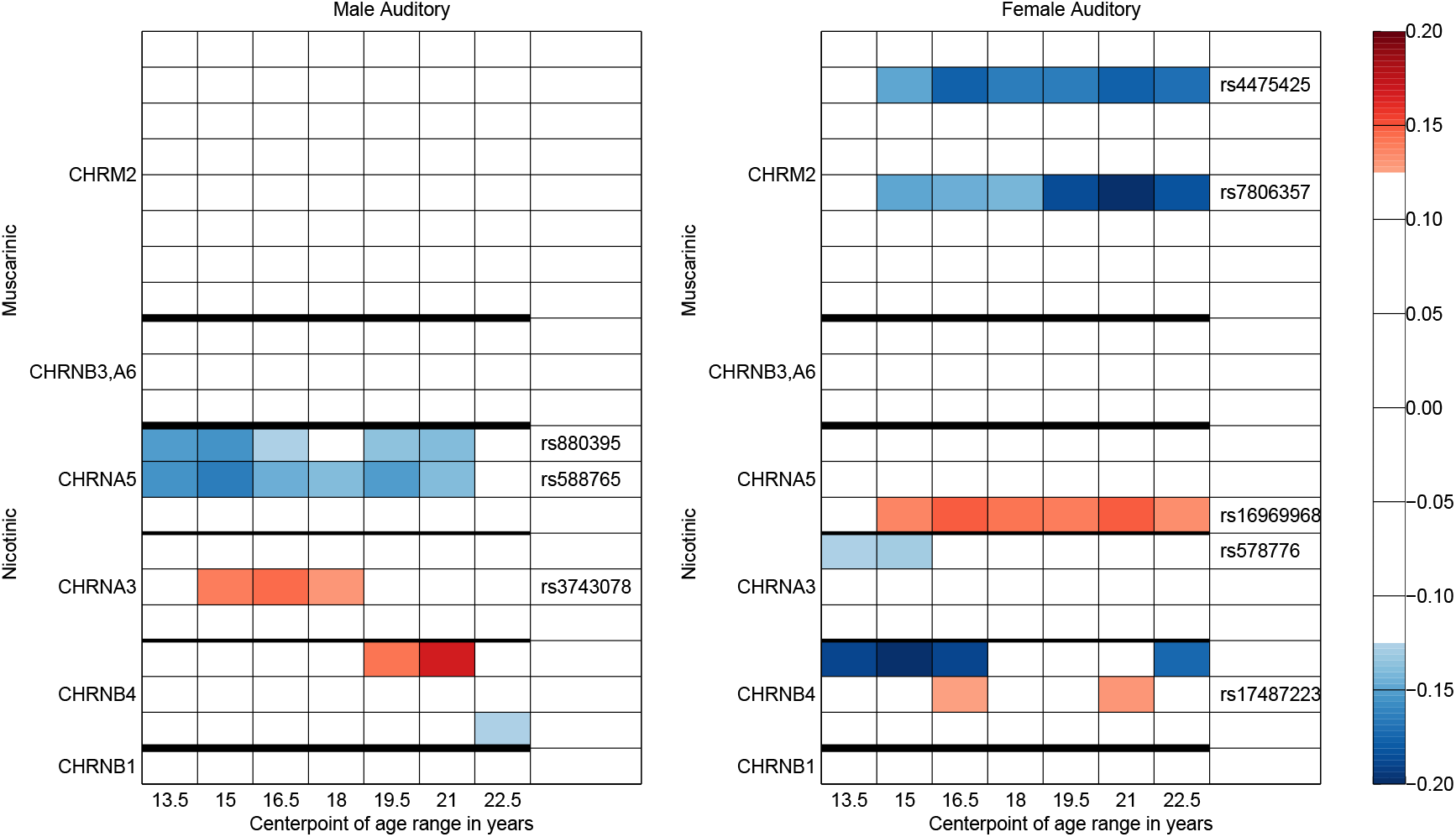
Identification of significant SNPs. Significant SNP-phenotype associations in the auditory modality. Color coding indicates signed value of significant median effect sizes over 3 year age ranges and all scalp locations.

Figure 4 shows significant association *differences* between males and females in each modality. Significant sex differences indicated in the left panel of this figure correspond approximately to the differences indicated between the left and right panels of Figure 3. The large sex differences found here are comparable to those exhibited in Cousminer et al. (2013) for human height in adolescents.

**Figure 4:**
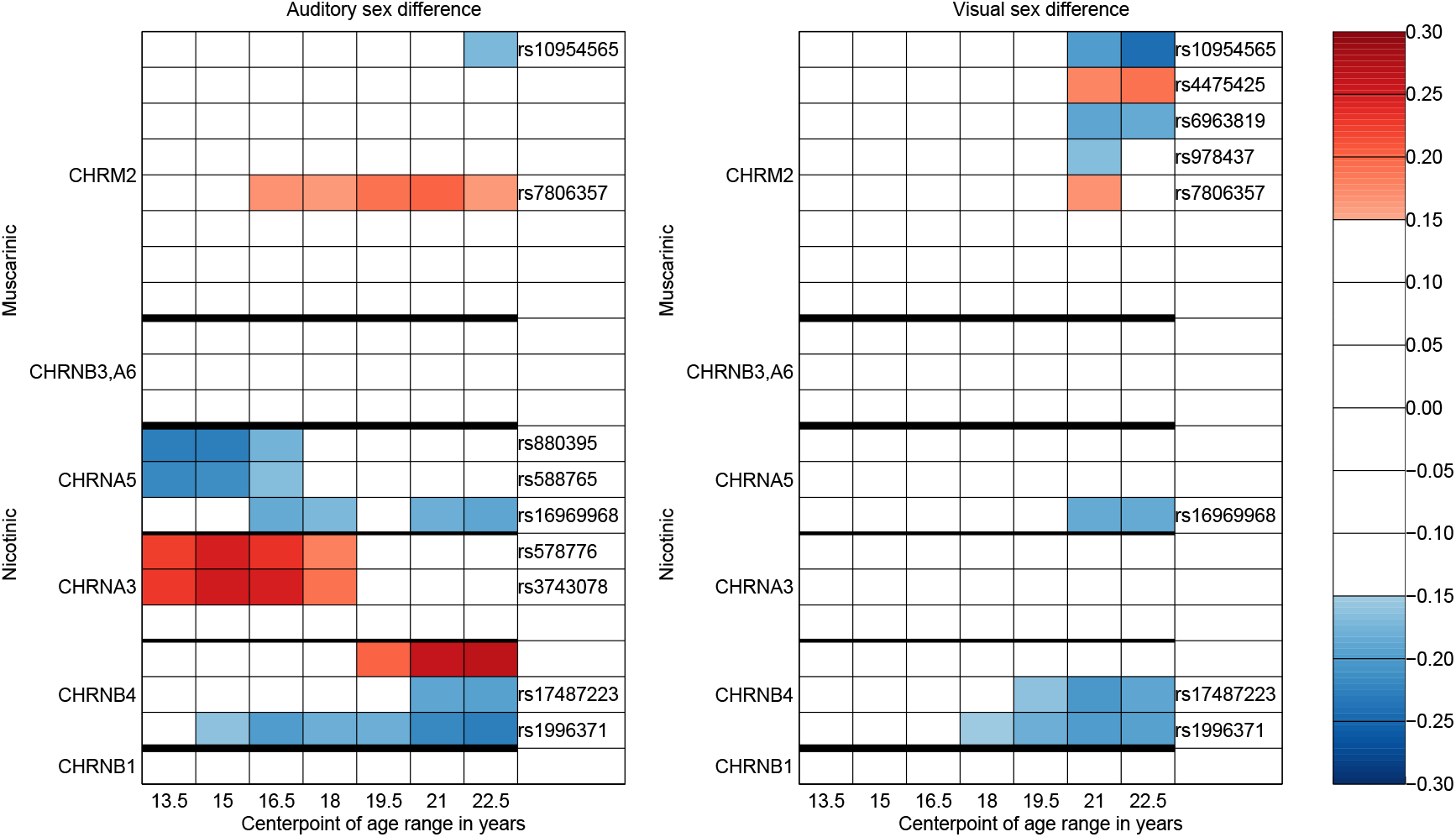
Significant sex differences in association. Significant SNP-phenotype male-female differences in associations in the auditory and visual modalities. Color coding indicates signed value of significantly different (male - female) median effect sizes over 3 year age ranges and all scalp locations.

## 4 Cholinergic - Theta ERO Associations: Systemic Results

### 4.1 Methods

The covariance of genotype-phenotype associations was determined by applying sparse principal component analysis (sPCA) to the genotypic profiles (vector of association effect sizes of each SNP) of all 36 phenotypes (9 electrodes × 2 stimulus modalities × 2 sexes) for each age range. Using sPCA yields a small set of SNPs which account for the bulk of the difference in genotypic profiles between the different phenotypes at each age range. The first two components of the sPCA were sufficient to represent the phenotypic variance between both stimulus modalities and sexes with respect to their pattern of genotypic associations. Bootstrapping confirmed the reliability of these results.

### 4.2 Results

Figure 5 illustrates the SNPs which are selected by this method. Note the similarity between this figure and figure 4, showing the concordance between SNP selection by *p*-value and by importance in separating phenotype groups.

**Figure 5:**
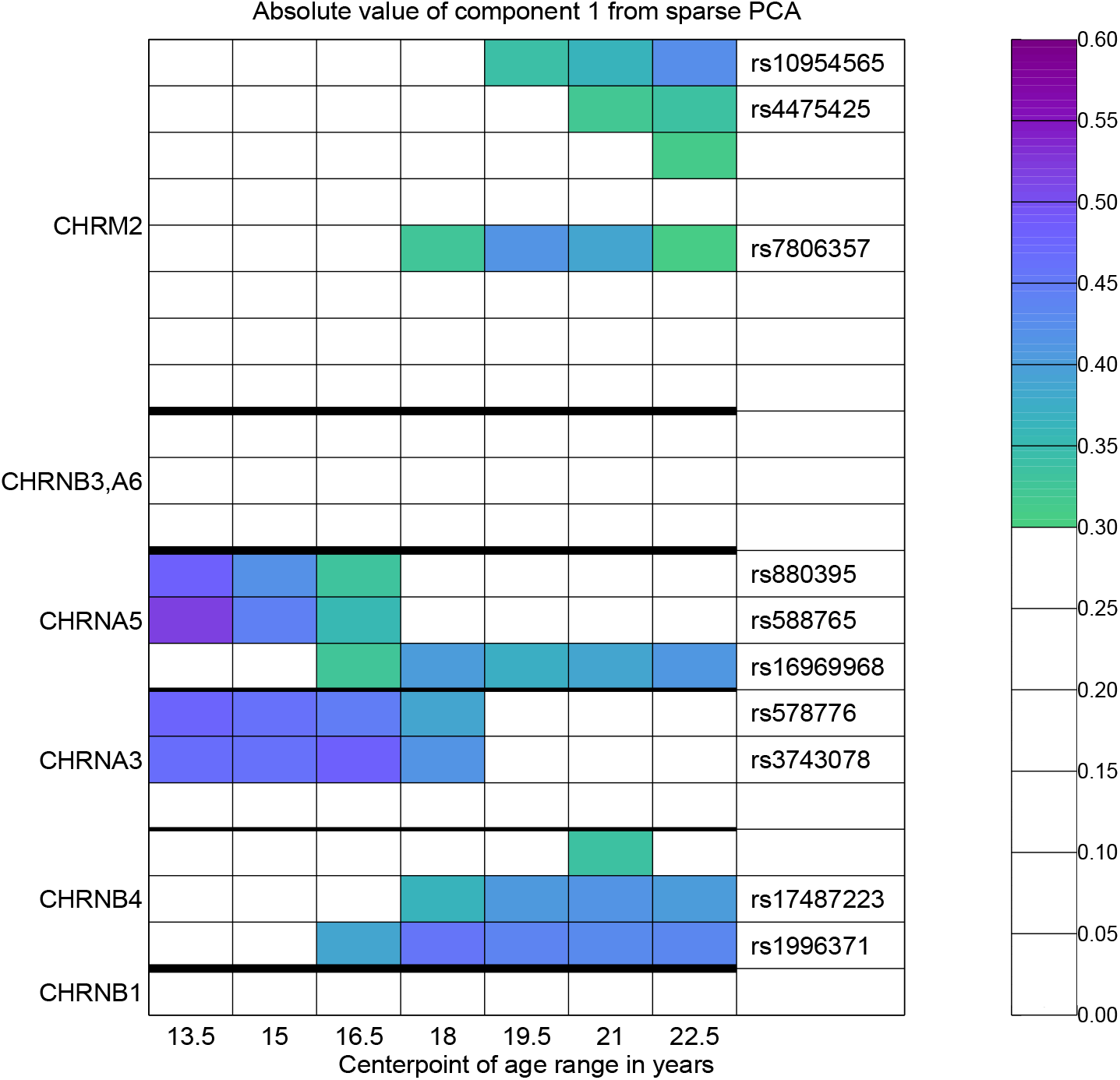
Identification of significant SNPs from the covariance structure of the genotype-phenotype associations. SNPs associated with phenotypic variance determined by the first component from age-specific sparse PCA. The sparsity constraint limits number of SNPs to parallel control by FDR threshold.

As shown in this figure, there appears to be a change in the set of SNPs associated with phenotypic variance over the course of development. This suggests there is a difference between those genetic factors which have a primary role in the timing of the development of the phenotype and those which have a primary role in phenotypic function.

Figure 6 illustrates the configuration of the phenotypes in the space defined by the first two principal components of SNP profiles from age-specific sparse PCA. Only the relation between the 4 groups in each plot can be compared between plots; as noted in the caption, the axes differ for each plot. Between sex differences for each modality are larger than between modality differences for each sex. How the spatial configurations of the phenotypes in the genotypic axes reflects the actual scalp topographies was not investigated.

**Figure 6:**
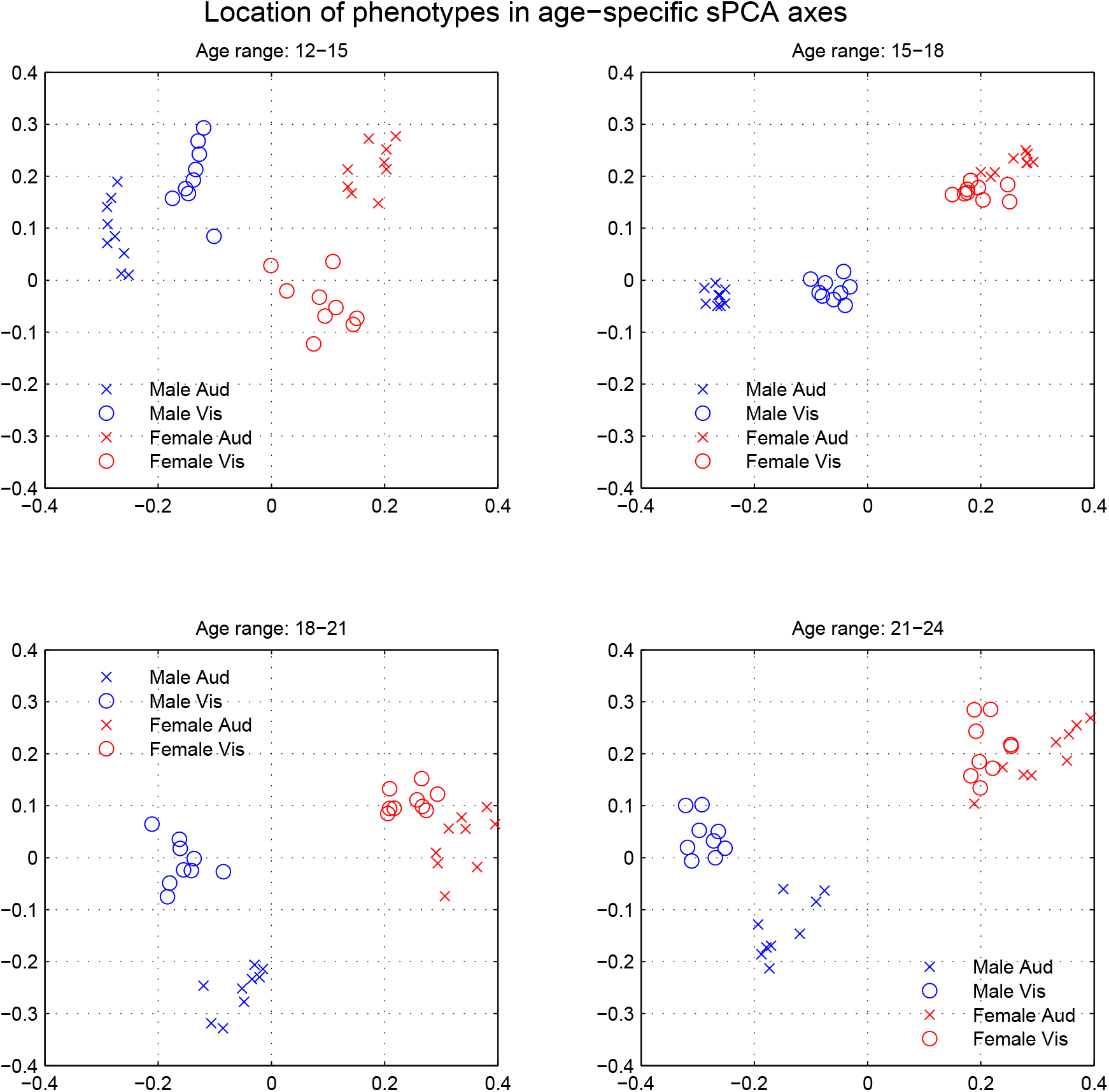
Discrimination of phenotypes by genotypic profile. Age specific phenotype-sex pairs for four age ranges in the space defined by the first two principal components of SNP profiles from age-specific sparse PCA. The positions of the phenotypes are particular to the individual plots, since the axes differ for each plot.

## 5 Research Directions

This study is the continuation of previously published work on the developmental trajectories of the theta EROs (Chorlian et al., 2015) and on the developmental association trajectories of theta EROs and SNPs from *KCNJ6* (Chorlian et al., 2017). The methods and results presented here will lead to future studies whose primary objectives will be:

- To determine the age variation in the genotypic factors influencing the developmental course of the phenotypes of interest.
- To identify and characterize the genetic systems contributing to that development by the analysis of temporal and phenotypic co-activation patterns.
- To characterize phenotypic systems by utilizing both phenotypic covariance and the covariance of genotypic profiles of phenotypes.
- To create appropriate measures for the quantification of genetic change in the developmental process.

The results suggest that candidate gene studies could be expanded to candidate system studies to yield considerable new information regarding the effect of various physiological systems in neurocognitive development. Additionally, the strong sex differences in the results suggest that future genetic analyses should incorporate sex × genotype interactions.

## 6 Data collection and analysis

This section summarizes the material in Chorlian et al. (2017), as this study uses the same data set used therein.

### Subjects

The sample comprised 2170 adolescents and young adults from the Prospective Study of the Collaborative Study on the Genetics of Alcoholism (COGA), a multisite collaboration (Begleiter et al., 1995), examined within the age range of 12 to 25 years.

### Genotyping

Genotyping was performed at Washington University School of Medicine in St. Louis on an OpenArray platform, and at Indiana University School of Medicine in Indianapolis on the Sequenom MassArray system.

### Electrophysiology

One important indicator of neurocognitive function is the P3 (or P300) response, evidenced in by the production of a large positive waveform with a peak between 300 ms. and 700 ms. after the presentation of a target stimulus. The P3 response is elicited by infrequently presented target stimuli in a stream of more frequently occurring non-target stimuli in auditory and visual target detection tasks. Frequency domain analysis of EEG recording suggests that the theta band event related oscillation (ERO) is a major constituent of the P3 response. Two target detection tasks, one visual, the other auditory, were used for this study. The response was recorded from frontal, central and parietal scalp locations. All six collaborating sites used identical experimental procedures and EEG acquisition hardware and software. Electrical activity was amplified 10,000 times using Neuroscan amplifiers and was recorded continuously over a bandwidth of 0.02-100.0 Hz on a Neuroscan system. Estimates of EROs, the localized power of non-stationary evoked potential time series, were obtained using the S-transform, a time-frequency representation method.

## 7 Retrospective view

The above sections represent the work of the authors as of early 2017. Since that time our thinking has evolved. The limitations of the above sections are clear:

- The trajectory analysis as presented is left at the level of individual SNP-phenotype trajectories.
- Isolated structural results were presented at separate age ranges.
- No attempt was made to characterize structural trajectories, either by direct analysis of the effect size matrices, or through analysis of the sPCA results both in the phenotypic and genotypic dimensions.
- Methods for structural analysis involve aggregation over phenotypes and over genotypes to create new time series with relatively easily interpretable results, and use of pairwise between-age distances of multivariate time series to create trajectories in a more abstract space through dimension reduction procedures.

### Statistical Supplement

The following material is derived from documents which were antecedents of the presentation text; the methodology primarily from Chorlian et al. (2017).

## 8 Statistical Methodology for the Analysis of Association Trajectories

### 8.1 Introduction

This study integrates the methods from both gene association studies and gene expression studies to study association trajectories. The study by Stead et al. (2006) of brain development in the male rat provides an exemplary model for comparison. The primary correspondence is between the gene expression levels in Stead et al. (2006) and the effect sizes in this study. Although effect sizes are signed and gene expression levels always positive, in gene expression studies measures of the difference from the mean are often the feature of interest. Trajectories of gene expression are shown in Stead et al. (2006, Figure 5) and PCA plots in Stead et al. (2006, Figure 2). While a single PCA in Stead et al. (2006) was sufficient to represent the variation in gene expression in both age (along the x-axis) and region (along the y-axis), in this study the complexity of the association relations necessitated the use of age specific PCAs, as can be observed in Figure 6 of the main document. The specific statistical methods described here are meant to deal with the situation of using a relatively small number of candidate SNPs with a possibly large age and phenotypic variation in effect sizes and phenotypes with a wide but overwhelmingly positive range of correlations.

### 8.2 Analysis of Association Trajectories

Extending a previous study of the development of theta band EROs (Chorlian et al., 2015) and as used in Chorlian et al. (2017), the model includes a genotypic value and a sex × genotype interaction, consistent with other studies of human development (Widén et al., 2010; Cousminer et al., 2013). Weights for individuals in the local linear regression model were adjusted to account for multiple observations on single individuals and co-presence of sibs in each of the regression calculations. The bandwidth for the kernel (Epanechnikov) was taken as 0.6 to minimize the mean squared error, although the variation in the size of the error was less than 2% over the range of bandwidths from 0.4 to 0.8.

In this study, the primary object of the analyses are the effect sizes of the SNPs obtained from the non-parametric regression calculations, although the *p*-values associated with these SNPs are also considered. The use of 21 SNPs and 18 phenotypes with separate effect sizes for males and females at 131 age centers produces 99036 effect size values. Effect sizes, unlike *p*-values, provide signed values which enable the direction of effect to be analyzed, and thus to more precisely characterize SNPs which have sex-specific and modality-specific effects. Effect sizes also are appropriately signed and scaled for covariance estimation, a key part of the analysis. The use of effect sizes also enables comparison between the results of this study and other studies which use different methodologies and have different sample sizes. In addition, arithmetic operations on effect sizes are easily interpreted, which is not the case with *p*-values.

As there are 5912 observations spread over a 13 year age-range, there is sufficient data to provide estimates of the means of the variables at one-tenth year intervals and to provide estimates of the mean rates of change of the variables as well. Since 90% of the successive observations of individuals have an interval of greater than 1.75 years, no longitudinal modeling at the shorter time scale (6 months) at which significant changes in the measured variables can occur is possible. (The estimate for the time scale is derived from Sullivan et al. (2011)). Results are not independent between models for different ages since the data in each of the 131 regression models has considerable overlap with the data used in the models for nearby ages. Significance levels and effect sizes were obtained from the regression calculations and corroborated using a non-parametric bootstrap method with 1000 resamplings. The bootstrapping process is described in Chorlian et al. (2017, p. 28). The regression method used in the calculation of the genotypic effect on the trajectories has the power to reliably detect effect sizes of greater than 0.1 based on the large number of subjects and observations included in the calculations of each parameter. Effect sizes of the major allele of significant SNPs typical of GWAS results in Kang et al. (2012) are of absolute value between 0.1 and 0.2.

### 8.3 Identification of significant SNPs

To characterize the age specific variation of genotypic effects on the phenotypic variables, significant SNPs are identified on an SNP specific basis. Two complementary methods are used, estimation of the false discovery rate using the method outlined in Storey and Tibshirani (2003), and the assessment of the concentration of effect sizes in particular SNPs. Although SNPs were selected on the basis of a likely involvement in the developmental process, it is not expected that all SNPs will be equally significant. Rather it is expected that significant effect sizes are concentrated in a relatively small number of SNP-phenotype combinations, since phenotypes are highly correlated across scalp electrodes for each sex-modality-age combination. Lack of concentration would suggest many of the large effect sizes (corresponding to low *p*-values) are the result of large random variation of effects across age and phenotype, and do not represent biologically stable phenomena. Concentration of effect is established by permutation tests which compare the realized distribution of effect sizes across SNP-phenotype combinations with the distribution of effect sizes of hypothetical random distributions of the realized effect sizes.

#### 8.3.1 Significant SNPs identified by thresholding p-values by false discovery rate

The evaluation of the significance of the association of the individual SNPs with theta EROs is determined by the selection of individual *p*-values based on control of the false discovery rate. Given the fact that this is a candidate gene study, in which at least a small proportion of the tested variables are expected to be significant, the control of the false discovery rate by the characteristics of data is appropriate. The method used to determine the false discovery rate for any particular level of *p*-value must be capable of dealing with correlated data, as *p*-values are correlated as a result of both the non-parametric regression method and the phenotypic correlations described in Chorlian et al. (2015). The method used is derived from that in Storey and Tibshirani (2003), which uses the distribution of the calculated *p*-values to determine the prevalence of results which reflect the absence of association, and correspondingly, the prevalence of results which reflect the presence of association.

Specifically, the distribution of *p*-values from non-associated SNP-phenotype pairs should be uniform on the interval from 0 to 1, while the the distribution of *p*-values from associated SNP-phenotype pairs should be concentrated near 0. From the estimation of the occurrence of non-associated SNP-phenotype pairs, it is possible to estimate the occurrence of associated SNP-phenotype pairs, and determine the false discovery rate, sometimes called a *q*-value, as a function of thresholding at different *p*-values. As with any thresholding technique, choice of a threshold is arbitrary, and the correspondence between *p*-values and *q*-values for levels *q* < .05 and *q* < .01 are determined. This procedure is appropriate for the evaluation of all obtained *p*-values collectively, as the algorithm deals with correlated values appropriately. This provides information on a fine-grained time scale. (The part of the original algorithm in Storey and Tibshirani (2003) which estimates the proportion of the non-associated results was modified to deal more robustly with truncated *p*-value distributions when necessary, and provides a somewhat more conservative estimate than does the original).

In the interest of condensation of information and ease of interpretation, as well as the expectation that meaningful biological processes must extend over more than the length of a year, results are provided for 7 overlapping 3 year age ranges extending from age 12 to 24. (Ranges of years of age: 12-15, 13.5-16.5, 15-18, 16.5-19.5, 18-21, 19.5-22.5, 21-24). Medians of *p*-values across the age-ranges and the four phenotype groups were calculated to provide coarsegrained results. The concentration of measure of the range of *p*-values introduced by use of the median was considerable, particularly in the range 0 < *p* < 10^− 2^ and introduced a corresponding bias in the direct application of the Storey-Tibshirani algorithm. As a consequence, the false discovery *proportion* threshold derived from the calculation on the raw *p*-values was used to determine the appropriate threshold for the control of the false discovery rate in the median *p*-values. A similar strategy is used to identify SNPs which had significant *differential* effect between sexes.

#### 8.3.2 Significant SNPs identified by concentration of effect sizes

To determine whether large effect sizes were distributed randomly across SNPs for sets of correlated phenotypes or concentrated in relatively few SNPs, the distribution of observed concentrations were tested against the hypothesis that there were no SNP-specific concentrations by a permutation test. The emphasis on SNPs is achieved by aggregating values across locations on the scalp, since phenotypic values are highly correlated over scalp locations.

As was done for the *p*-values, to simplify the analysis of age variation in the association trajectories, the absolute values of the 131 age-specific effect sizes for each SNP-phenotype pairing were averaged into 7 age-ranges of three years duration overlapping by one and one half years covering ages 12-24. The same procedure was applied to the absolute value of the difference between effect sizes for males and females for corresponding modality-locality-SNP combinations, The averaging procedure was in part adopted to ensure that the values represented biologically meaningful phenomena and were not the result of random short-term fluctuations in the data.

In order to identify the significant SNPs, the values of the age-range averaged absolute effect sizes for each SNP were averaged over the 9 electrodes for each of the 4 sex-modality combinations, resulting in 4 averages from each of the seven age ranges for each SNP for the observed data. This provides 147 (21 SNPs × 7 age-ranges) SNP-age range effect sizes for each of the 4 sex-modality combinations. Given the generally high correlation between the theta EROs at the 9 electrodes (all correlations greater than 0.5, fewer than 3% less than 0.6), this is an additional method of reducing the effect of spurious large effect sizes on the identification of significant SNPs.

Permutation tests were carried out to test the null hypothesis that the distribution of the averaged absolute value of effect size values was not SNP-specific, that is, that a random distribution of the observed effect sizes among SNPs could account for the observed variability of effect sizes among SNP. To characterize the null hypothesis, the absolute values of the effect sizes from different SNPs were used when the averaging over the electrodes to create values for “pseudo”-SNPs in a manner similar to the averaging over the actual SNPs described in the previous paragraph. Since the data contains 21 SNPs, 21 pseudo-SNPs were created for each permutation calculation. For each pseudo-SNP, the average of the effect sizes over the 9 electrodes is not from a single SNP but from 9 different SNPs, no combination of which is repeated among the 21 pseudo-SNPs. That is, 9 distinct permutations of the effect sizes of the 21 SNPs are produced, one for each electrode, resulting in a matrix of 9 × 21 values for each sex-modality-age combination. Then the effect sizes are averaged across electrodes for each of the 21 pseudo-SNPs created by the permutation process. This is done for each sex-modality combination, resulting in 4 averages from each of the seven age ranges for each pseudo-SNP for the observed data. Randomly permuting the SNPs contributing to the averaging across the electrodes ensures that the effect sizes from all the SNPs are used. Each permutation procedure instance produces the same number of random groups of values for pseudo-SNPs as is produced for the actual SNPs. No permutations of age or phenotype are performed in order to preserve the covariance of those items from the original calculation. A large number (1000) of permutation sets were calculated, and for each of the seven age ranges, the mean and standard deviations over all the pseudo-SNPs in each of the permutation procedures are calculated to provide the statistical properties associated with the null hypothesis. It should be noted that the permutation test results for any one SNP-age-sex-phenotype combination, unlike the actual effect sizes, is a function of the effect sizes of all the other SNPs for that age-sex-phenotype combination, and thus reflect medium scale characteristics of the genotypic effects of the cholinergic system SNPs. For each of the actual SNPs the difference between the means over age ranges of the mean effect sizes of SNPs and the group mean of the pseudo-SNPs for the corresponding age ranges measured in standard deviations of the pseudo-SNPs values is obtained. The significance of the value for the actual SNPs is evaluated by a z-test.

The permutation tests are scale-free, and their use to identify important SNPs depends upon the absolute values of the effect sizes themselves. Thus some effect size thresholding is necessary to make this method useful for identifying significant SNPs; conversely, if all effect sizes are significantly large, then the method is superfluous. The averaging procedure serves to reduce the variance of the aggregated data, and combined with information from bootstrapping, ensures the reliability of the effect size estimates, whose importance can be gauged from the results of other genetic studies. We note that effect sizes of the major allele of significant SNPs typical of GWAS results in Kang et al. (2012) are of absolute value between 0.1 and 0.2. In the particular distribution of effect sizes obtained in this study, the method confirmed, that except for a single SNP, a threshold of .15 in effect size would correspond to a threshold of 10^− 4^ in p-value, as shown in the top panel of Figure 7.

**Figure 7:**
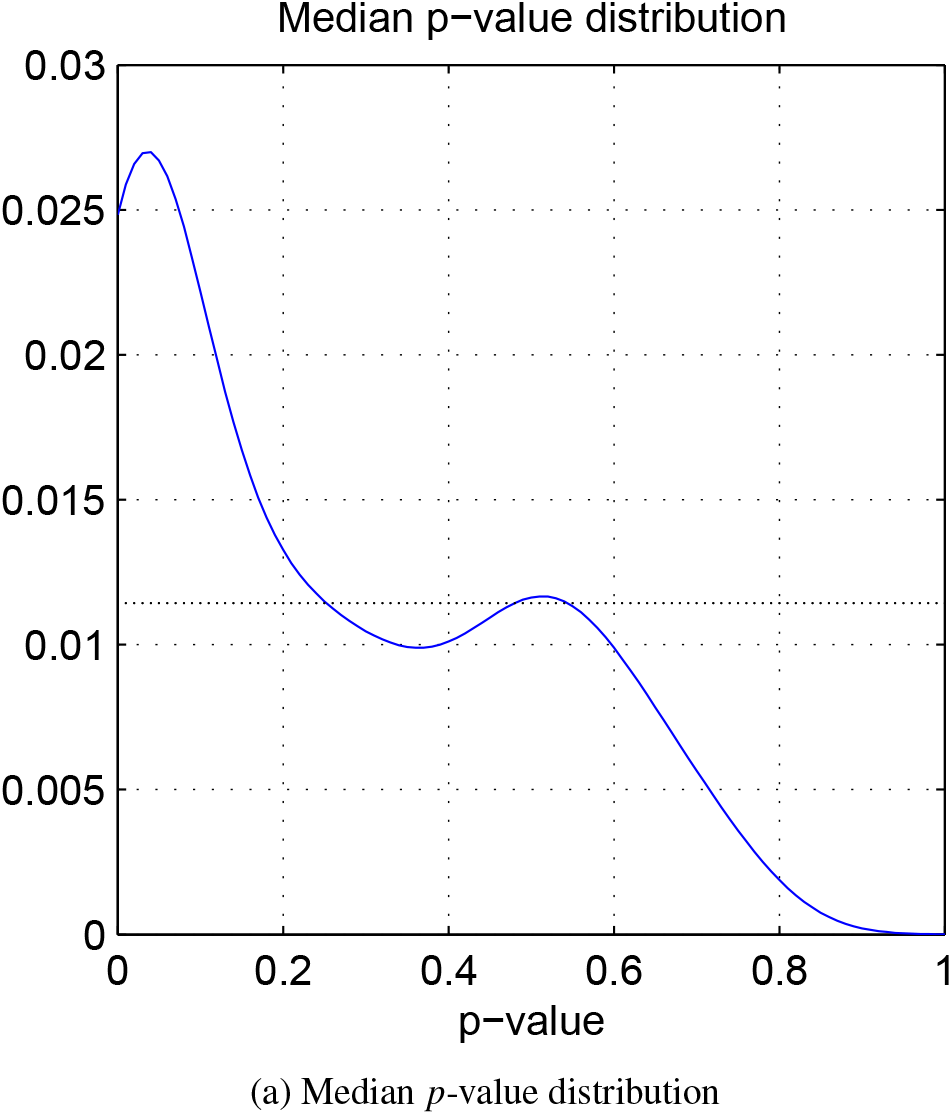
Median *p*-value.

Absolute values of effect sizes follow a folded normal distribution with variance 0.0025 corresponding to a normal distribution with variance 0.0069. However mean effect sizes averaged across correlated phenotypes and 3 year age ranges are neither uniformly nor normally distributed across SNPs as shown by the permutation test results. The distribution of the permutation test results, expressed in terms of z-scores, are not symmetrical but have a peak at about -2 standard deviations and a long rightward tail beginning about +2 standard deviations above the mean as shown in Figure 8. The fraction of effect sizes less than .1 among the SNP-age-phenotype combinations selected by having permutation scores greater than the inverse normal cdf determined by *p*-values in the range .05 to 10^− 4^ was linearly related to the logarithm of the p-value, decreasing from about 30% for *p* = .05 to 15% for *p* = 10^− 4^. For *p* < 10^− 4^, the rate of decrease diminished from the rate over the range of .05 to 10^− 4^. Further observation of the distribution suggests that the SNP-age combinations fall into three groups, non-significant, null, and significant.

**Figure 8:**
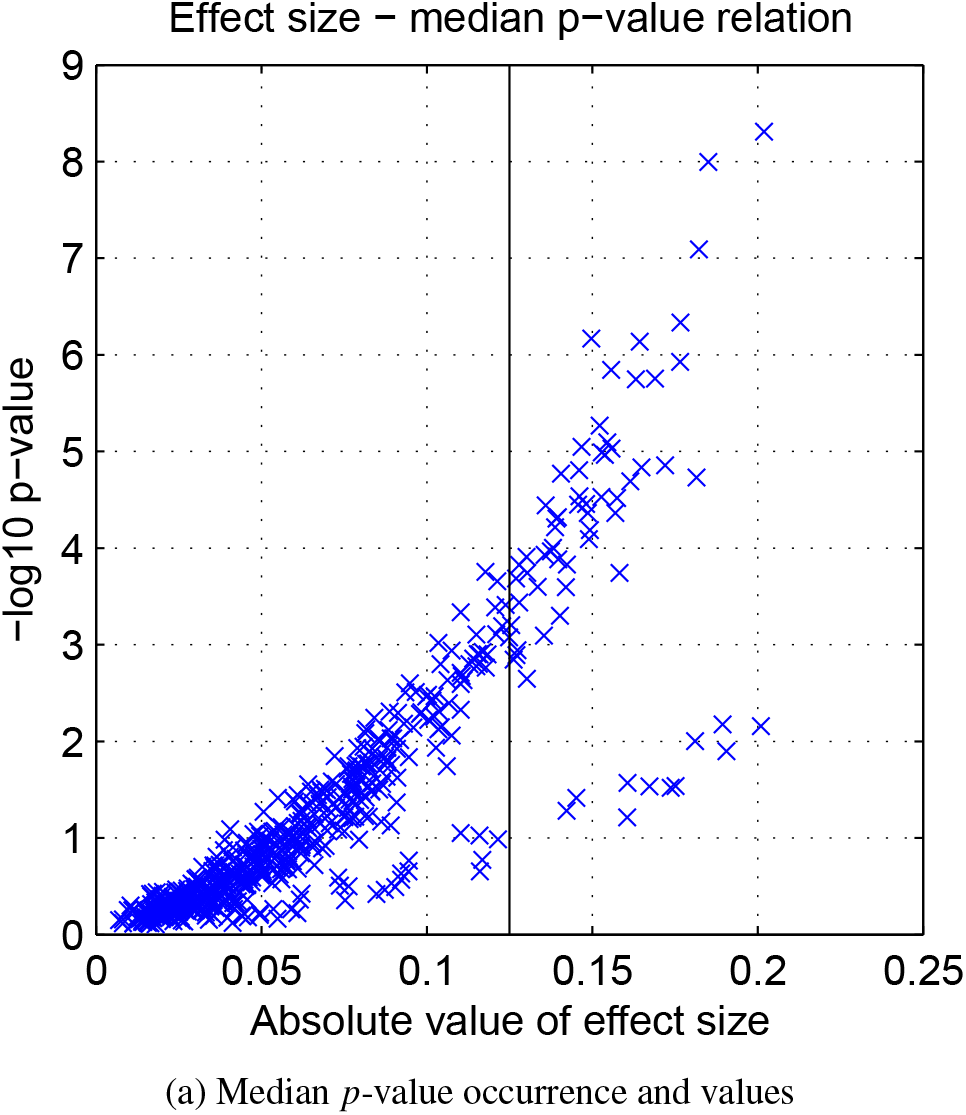
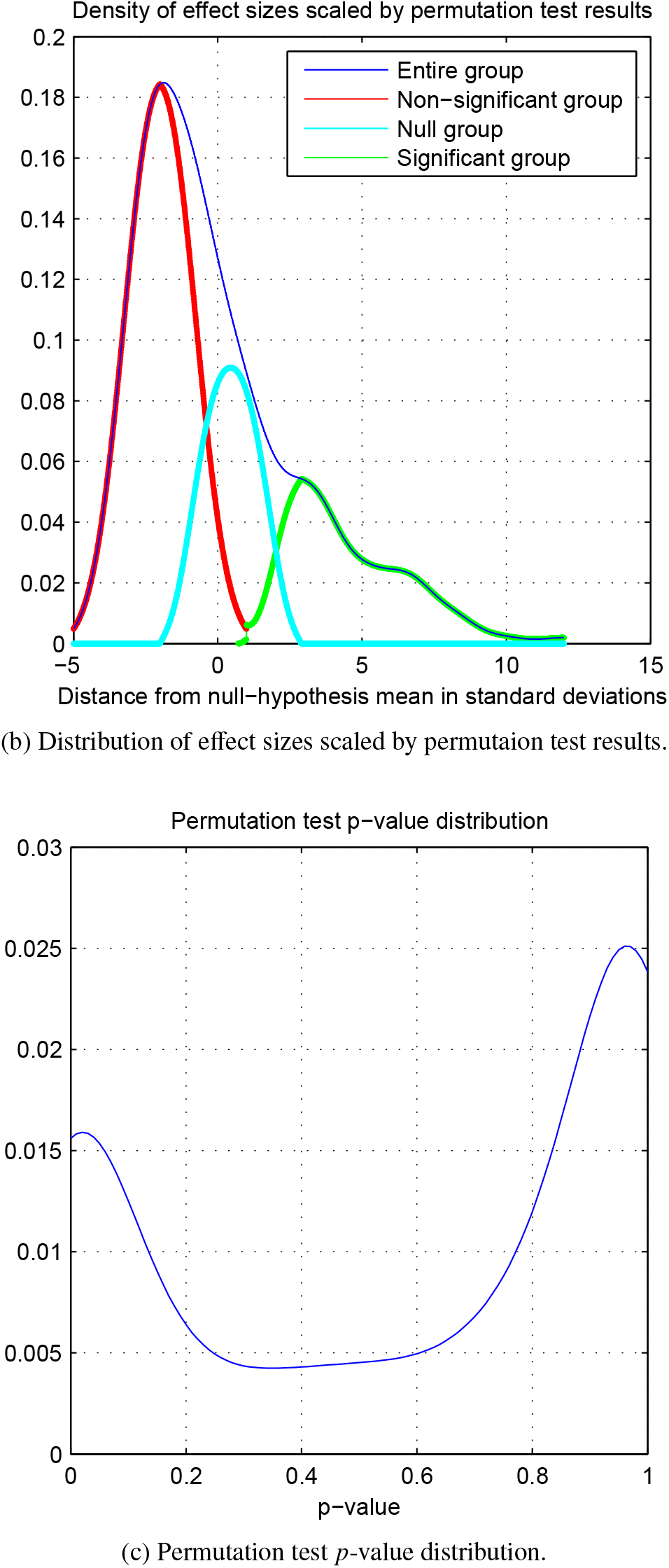
Permutation test results.

Any selection of SNP-age-phenotype combinations by thresholding of permutation scores will result in effect sizes specific to individual phenotypes which are less than 0.1 (or comparable value) being included in the select group, and effect sizes which are greater than 0.1 excluded from the select group. The exclusions are justified on the basis that they represent either sporadic occurrences over age and phenotype and therefore not of biological interest or else false positives; the inclusions reflect genetic heterogeneity in scalp localization in which large effect sizes in one region will counterbalance smaller effect sizes in other regions. Determination of the significance level for the *p*-values associated with the SNP-age combinations from the permutation was made by adapting the method of Storey and Tibshirani (2003) to determine the *q*-value (false discovery rate associated with the particular *p*-value), and establish the false discovery rate corresponding, if somewhat indirectly, to setting a threshold for effect sizes.

#### 8.4 Identification of SNPs contributing to Phenotypic Variance

Principal component analysis was not applied to the data set corresponding to the entire age range because the use of PCA on the individual age ranges revealed that the first components varied considerably across the age ranges. Application of PCA to the entire age range would have identified components which were only remotely interpretable compared to the components identified in the individual age ranges. Instead the seven data sets consisting of the effect sizes for individual phenotypes as determined by the age range averaging described in section 8.3.2 were used in the analysis.

#### 8.5 Analysis of sPCA results

Sparse principal component analysis (sPCA) was used to determine genotypic profiles which discriminated between modalities and sexes. In contrast to the tests for significant SNPs, sPCA identifies only individual SNPs which have differential effects between sexes, modalities, or localities. Parameters for sparseness were two components with no more than 10 SNPs per component applied to the aggregated effect sizes described in section 8.3.2. Given the restriction of 10 SNPs in each component, two-thirds of the 147 SNP-age range combinations never appear in the first component. This is the non-significant group for determining the empirical null distribution. The significance of the members of the group non-zero components are evaluated based on the empirical null distribution. The same sPCA analysis was applied to the 1000 bootstrap samples described in section 8.2 and the proportion of occurrence of each of the 147 SNP-age range combinations in the first sPCA component of the 1000 bootstrap samples determined, as well as the frequency of occurrence of the significant and non-significant SNP-age range combinations from the actual sample. The 95th percentile of cumulative distribution function of the fraction of the occurrences of each of the members of the 99 members non-significant group in the sparse PCA across the bootstrap sample acts as the criterion for significance at the .05 level; every snp-age range combination which occurs in a greater fraction of bootstrap instances than that of the significance criterion is found to be significant. This is an adaptation of the method described in Efron (2008).

**Figure 9:**
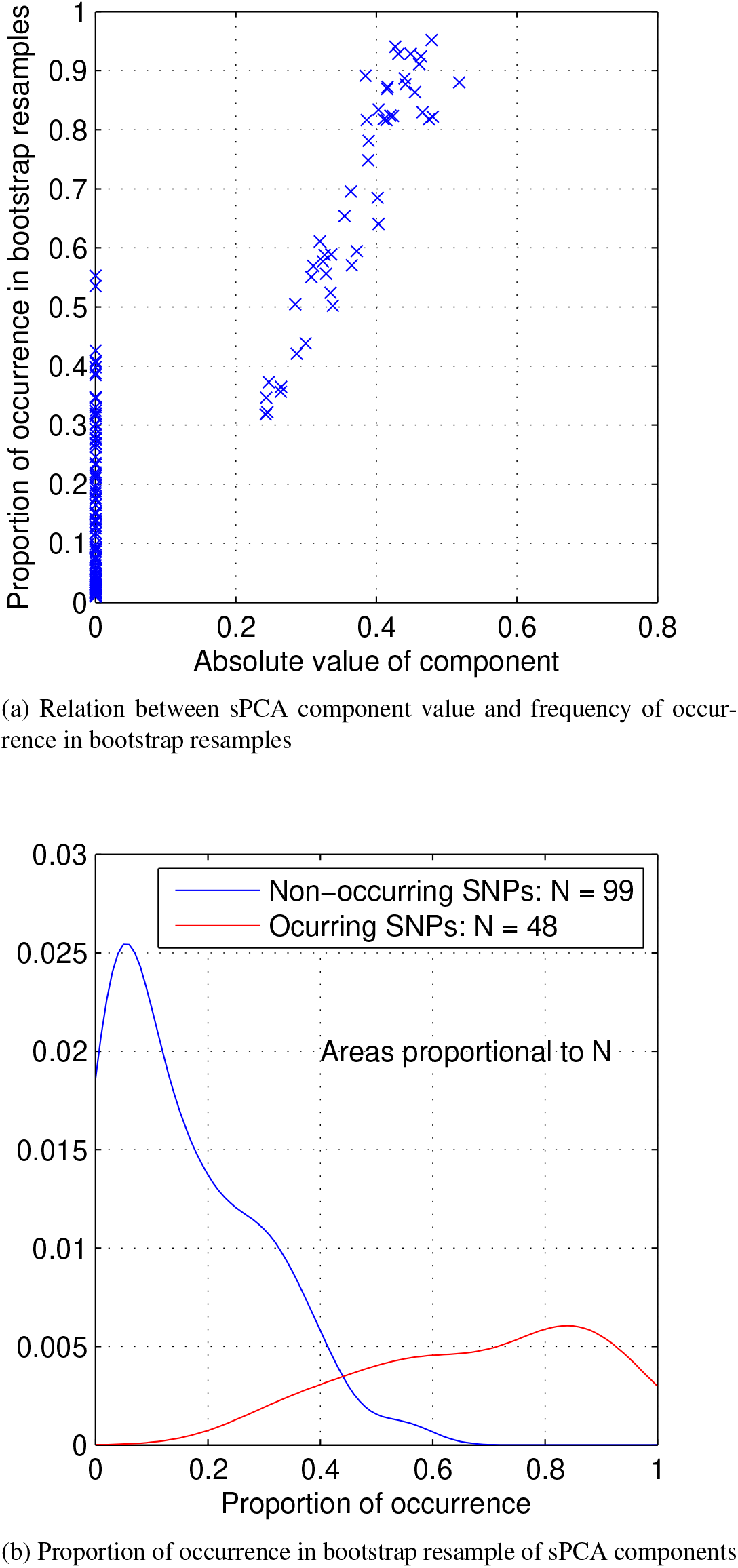
Sparse PCA comparison with bootstrap frequency of occurance.

#### 8.6 Retrospective view

Further reflection on the these topics suggest that a multiscale approach, that is, with the aggregation process taking place within each of the four phenotypic classes, would provide useful additional information. Correspondingly, larger studies with greater numbers of phenotypic and genotypic classes could have several different levels of scaling. Exactly how this would be implemented is still to be determined.

## 9 Acknowledgements

**As of 2017** The Collaborative Study on the Genetics of Alcoholism (COGA), Principal Investigators B. Porjesz, V. Hesselbrock, H. Edenberg, L. Bierut, includes eleven different centers: University of Connecticut (V. Hesselbrock); Indiana University (H.J. Edenberg, J. Nurnberger Jr., T. Foroud); University of Iowa (S. Kuperman, J. Kramer); SUNY Downstate (B. Porjesz); Washington University in St. Louis (L. Bierut, J. Rice, K. Bucholz, A. Agrawal); University of California at San Diego (M. Schuckit); Rutgers University (J. Tischfield, A. Brooks); University of Texas Health Science Center at San Antonio (L. Almasy), Virginia Commonwealth University (D. Dick), Icahn School of Medicine at Mount Sinai (A. Goate), and Howard University (R. Taylor). Other COGA collaborators include: L. Bauer (University of Connecticut); D. Koller, J. McClintick, S. O’Connor, L. Wetherill, X. Xuei, Y. Liu (Indiana University); G. Chan (University of Iowa; University of Connecticut); D. Chorlian, N. Manz, C. Kamarajan, A. Pandey (SUNY Downstate); J.-C. Wang, M. Kapoor (Icahn School of Medicine at Mount Sinai) and F. Aliev (Virginia Commonwealth University). A. Parsian and M. Reilly are the NIAAA Staff Collaborators.

**As of 2023** The Collaborative Study on the Genetics of Alcoholism (COGA), Principal Investigators B. Porjesz, V. Hesselbrock, T. Foroud; Scientific Director, A. Agrawal; Translational Director, D. Dick, includes ten different centers: University of Connecticut (V. Hesselbrock); Indiana University (H.J. Edenberg, T. Foroud, Y. Liu, M.H. Plawecki); University of Iowa Carver College of Medicine (S. Kuperman, J. Kramer); SUNY Downstate Health Sciences University (B. Porjesz, J. Meyers, C. Kamarajan, A. Pandey); Washington University in St. Louis (L. Bierut, J. Rice, K. Bucholz, A. Agrawal); University of California at San Diego (M. Schuckit); Rutgers University (J. Tischfield, D. Dick, R. Hart, J. Salvatore); The Children’s Hospital of Philadelphia, University of Pennsylvania (L. Almasy); Icahn School of Medicine at Mount Sinai (A. Goate, P. Slesinger); and Howard University (D. Scott). Other COGA collaborators include: L. Bauer (University of Connecticut); J. Nurnberger Jr., L. Wetherill, X., Xuei, D. Lai, S. O’Connor, (Indiana University); G. Chan (University of Iowa; University of Connecticut); D.B. Chorlian, J. Zhang, P. Barr, S. Kinreich, G. Pandey (SUNY Downstate); N. Mullins (Icahn School of Medicine at Mount Sinai); A. Anokhin, S. Hartz, E. Johnson, V. McCutcheon, S. Saccone (Washington University); J. Moore, F. Aliev, Z. Pang, S. Kuo (Rutgers University); A. Merikangas (The Children’s Hospital of Philadelphia and University of Pennsylvania); H. Chin and A. Parsian are the NIAAA Staff Collaborators.

We continue to be inspired by our memories of Henri Begleiter and Theodore Reich, founding PI and Co-PI of COGA, and also owe a debt of gratitude to other past organizers of COGA, including Ting-Kai Li, P. Michael Conneally, Raymond Crowe, and Wendy Reich, for their critical contributions. This national collaborative study is supported by NIH Grant U10AA008401 from the National Institute on Alcohol Abuse and Alcoholism (NIAAA) and the National Institute on Drug Abuse (NIDA).

## 10 Supplementary material

### 10.1 Table of Cholinergic SNPs

**Table 1:**
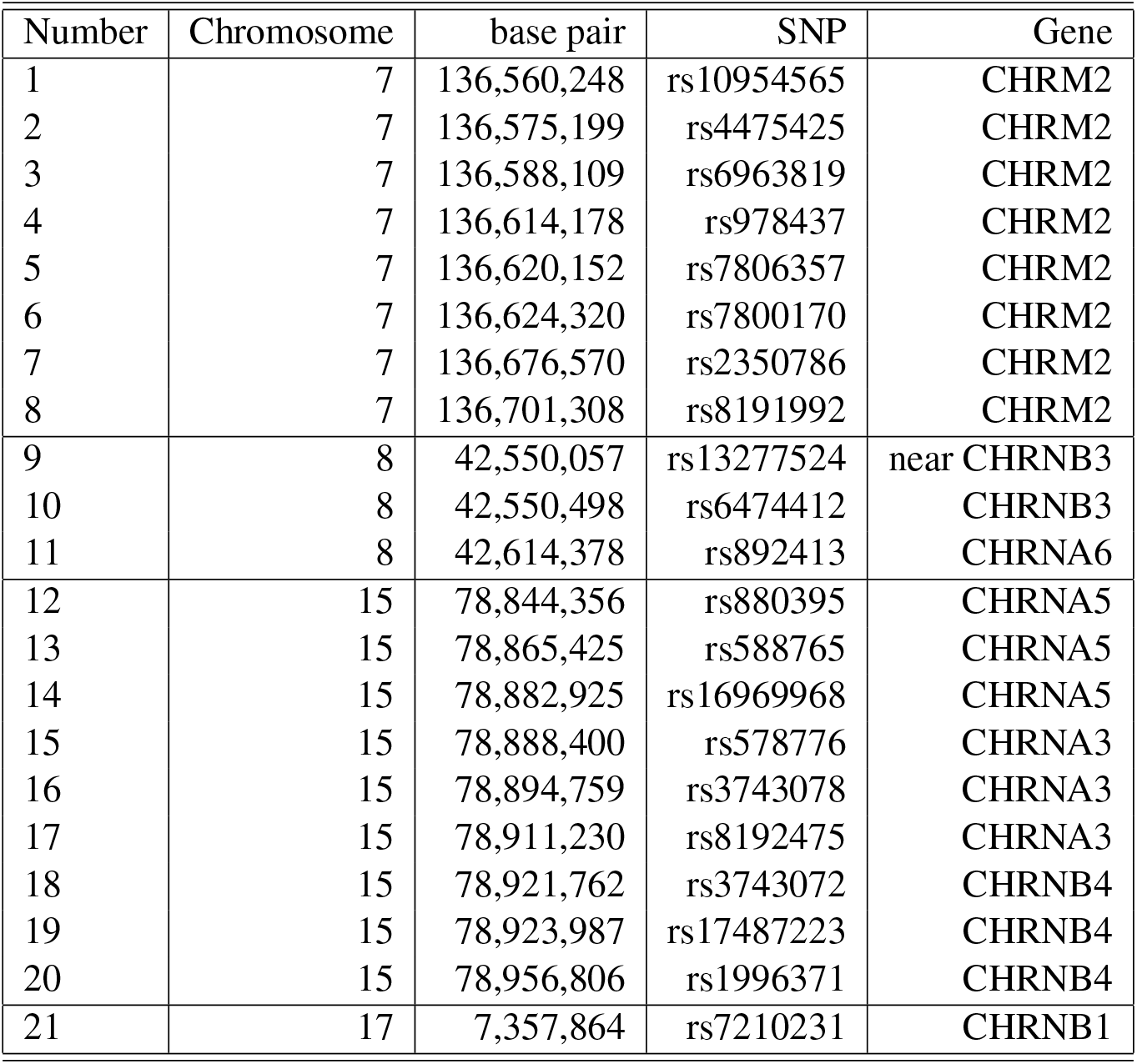
Cholinergic SNPs ordered by chromosome – base pair

## References

Begleiter, H., Reich, T., Hesselbrock, V.M., Porjesz, B., Li, T.K., Schuckit, M.A., et al., (1995) The Collaborative Study on the Genetics of Alcoholism. Alcohol Health Res. World 19:228–236.

Chorlian DB, Rangaswamy M, Manz N, Kamarajan C, Pandey AK, Edenberg H, Kuperman S, Porjesz B. (2015) Gender modulates the development of Theta Event Related Oscillations in Adolescents and Young Adults. Behav Brain Res. 292:342–52

Chorlian DB, Rangaswamy M, Manz N, Kamarajan C, Pandey AK, Wang JC, Wetherill L, Edenberg H, Porjesz B. (2017) Genetic correlates of the development of Theta Event Related Oscillations in Adolescents and Young Adults. Int J Psychophysiol. 2017 May;115:24–39.

Cousminer DL, Berry DJ, Timpson NJ, Ang W, Thiering E, Byrne EM, Taal HR, Huikari V, Bradfield JP, Kerkhof M, Groen-Blokhuis MM, Kreiner-M Kreiner-Møller E, Marinelli M, Holst C, Leinonen JT, Perry JR, Surakka I, Pietiläinen O, Kettunen J, Anttila V, Kaakinen M, Sovio U, Pouta A, Das S, Lagou V, Power C, Prokopenko I, Evans DM, Kemp JP, St Pourcain B, Ring S, Palotie A, Kajantie E, Osmond C, Lehtimäki T, Viikari JS, Kähönen M, Warrington NM, Lye SJ, Palmer LJ, Tiesler CM, Flexeder C, Montgomery GW, Medland SE, Hofman A, Hakonarson H, Guxens M, Bartels M, Salomaa V; ReproGen Consortium, Murabito JM, Kaprio J, Sørensen TI, Ballester F, Bisgaard H, Boomsma DI, Koppelman GH, Grant SF, Jaddoe VW, Martin NG, Heinrich J, Pennell CE, Raitakari OT, Eriksson JG, Smith GD, Hyppönen E, Järvelin MR, McCarthy MI, Ripatti S, Widén E; Early Growth Genetics (EGG) Consortium. (2013) Genome-wide association and longitudinal analyses reveal genetic loci linking pubertal height growth, pubertal timing and childhood adiposity. Hum Mol Genet. 22(13):2735–47.

Efron, B. (2008) Microarrays, empirical Bayes and the two-groups model. Stat Sci. 23(1):1:22

Jones KA, Porjesz B, Almasy L, Bierut L, Dick D, Goate A, Hinrichs A, Rice JP, Wang JC, Bauer LO, Crowe R, Foroud T, Hesselbrock V, Kuperman S, Nurnberger J Jr, O’Connor SJ, Rohrbaugh J, Schuckit MA, Tischfield J, Edenberg HJ, Begleiter H. (2006b) A cholinergic receptor gene (CHRM2) affects event-related oscillations. Behav Genet. 36(5):627–39.

Kang SJ, Rangaswamy M, Manz N, Wang JC, Wetherill L, Hinrichs T, Almasy L, Brooks A, Chorlian DB, Dick D, Hesselbrock V, Kramer J, Kuperman S, Nurnberger J Jr, Rice J, Schuckit M, Tischfield J, Bierut LJ, Edenberg HJ, Goate A, Foroud T, Porjesz B. (2012) Family-based genome-wide association study of frontal theta oscillations identifies potassium channel gene KCNJ6. Genes Brain Behav. 11(6):712–9.

Stead JD, Neal C, Meng F, Wang Y, Evans S, Vazquez DM, Watson SJ, Akil H. (2006) Transcriptional profiling of the developing rat brain reveals that the most dramatic regional differentiation in gene expression occurs postpartum. J Neurosci 26(1):345–53.

Storey JD, Tibshirani R. (2003) Statistical significance for genomewide studies. Proc Natl Acad Sci U S A. 100(16):9440–5.

Sullivan EV, Pfefferbaum A, Rohlfing T, Baker FC, Padilla ML, Colrain IM. (2011) Developmental change in regional brain structure over 7 months in early adolescence: comparison of approaches for longitudinal atlas-based parcellation. Neuroimage. 57(1):214–24.

Widén E, Ripatti S, Cousminer DL, Surakka I, Lappalainen T, Järvelin MR, Eriksson JG, Raitakari O, Salomaa V, Sovio U, Hartikainen AL, Pouta A, McCarthy MI, Osmond C, Kajantie E, Lehtimäki T, Viikari J, Kähönen M, Tyler-Smith C, Freimer N, Hirschhorn JN, Peltonen L, Palotie A. (2010) Distinct variants at LIN28B influence growth in height from birth to adulthood. Am J Hum Genet. 86(5):773–82.

